# Involvement of FKBP5, but not of stress, in alcohol memory reconsolidation

**DOI:** 10.1101/2023.12.12.570934

**Authors:** Nofar Rahamim, Coral Aronovici, Mirit Liran, Koral Goltseker, Matar Levin-Greenwald, Tim Heymann, Felix Hausch, Segev Barak

**Affiliations:** Sagol School of Neuroscience, Tel Aviv University, Tel Aviv 69978, Israel; School of Psychological Sciences, Tel Aviv University, Tel Aviv 69978, Israel; Faculty of Life Sciences, Department of Neurobiology, Tel Aviv University, Tel Aviv 69978, Israel; Zuckerman Mind Brain Behavior Institute, Columbia University, New York, NY 10027, USA; Department of Chemistry, Technical University Darmstadt, Darmstadt 64287, Germany; Department of Proteomics and Signal Transduction, Max Planck Institute of Biochemistry, Am Klopferspitz 18, Martinsried 82152, Germany; Centre for Synthetic Biology, Technical University of Darmstadt, Darmstadt 64283, Germany

**Keywords:** Alcohol, Addiction, Memory reconsolidation, Stress, HPA axis, FKBP5, FKBP51

## Abstract

Relapse is a fundamental challenge in drug addiction, often evoked by exposure to drug-associated cues. Upon retrieval, memories become temporarily labile before re-stabilizing in a process termed reconsolidation. Therefore, targeting the reconsolidation process offers a therapeutic approach for relapse prevention via the disruption of the drug-cue memories. We recently demonstrated that retrieval of contextual alcohol memories increased the expression of the mRNA encoding for FK506 binding protein 51 (FKBP51), a regulator of the hypothalamic-pituitary-adrenal (HPA) axis. Here, we explored the role of the HPA axis, and FKBP5/FKBP51 in particular, in the reconsolidation of alcohol memories. We found that the FKBP51 inhibitor SAFit2 given before alcohol-memory retrieval using contextual cues prevented the extinction of alcohol place preference behavior in female mice, suggesting that this protein may play a role in cognitive flexibility in a sex-dependent manner. Conversely, the retrieval of alcohol memories using an odor-taste cue did not affect *Fkbp5* expression in rats with a history of chronic alcohol consumption, suggesting that FKBP5 may play a differential role in different alcohol-associated memories. In addition, we provide evidence for HPA axis activation following alcohol memory retrieval, by showing that exposure to an alcohol-associated context led to elevated corticosterone secretion. However, we found that the reconsolidation process was unaffected by HPA axis-related manipulations, namely stress exposure, and administration of corticosterone or the glucocorticoid receptors inhibitor, mifepristone. Our results suggest that although FKBP5 can affect cognitive flexibility, and thereby impact the reconsolidation of alcohol memories, this effect is not likely mediated by HPA axis-related mechanisms.

## 1. Introduction

Alcohol use disorder (AUD) is a chronic psychiatric disorder ^1,2^ with limited available treatments ^3,4^, and more than half of the patients relapse within a year of abstinence ^5^. Relapse is often triggered by re-exposure to cues that had been associated with the reinforcing effect of alcohol and can elicit craving ^5–11^. Therefore, understanding the psychological and neurobiological basis of cue-alcohol memories can advance novel memory modulation methods to prevent relapse ^10–12^.

A promising approach for the disruption of maladaptive memories is to target the process of memory reconsolidation ^10,13,14^. Thus, it is believed that reactivation of stable memories via their retrieval initiates a 5-6-hour window, during which the memories become flexible and labile, before their re-stabilization, in the process of “memory reconsolidation” ^15,16^. Previous studies have shown that pharmacological or behavioral manipulations applied during the “reconsolidation window” can impair the reconsolidation of drug/alcohol-associated memories, and attenuate the subsequent expression of the targeted memory, i.e., relapse ^12,14,17–22^.

It is well-established that the process of memory reconsolidation requires *de-novo* protein synthesis ^20,23^, and it has been suggested that gene transcription is also required ^24,25^. In line with the latter, using RNA sequencing, we recently characterized increases in the mRNA expression of a set of genes in the hippocampus and medial prefrontal cortex (mPFC) of mice following the retrieval of alcohol memories ^25^. One of the genes exhibiting increased expression in the mPFC upon alcohol memory retrieval is *Fkbp5*, which encodes for FK506 binding protein 51 (FKBP51), an important regulator of the hypothalamic-pituitary-adrenal (HPA) axis stress response system ^26^. When bound to the glucocorticoid receptor (GR) complex, FKBP51 reduces the receptor’s affinity for cortisol/corticosterone ^26–28^. When cortisol is abundant upon activation of the HPA axis, FKBP51 dissociates from the complex, allowing GR translocation into the nucleus and the initiation of negative feedback on HPA axis activity ^27,29^. Within the nucleus, GR acts as a transcription factor, with *FKBP5* itself being one of its transcriptional targets ^27,30^. Indeed, activation of the HPA axis, triggered by stress ^31^ or by alcohol exposure ^32–34^, increases brain *FKBP5* expression.

Importantly, frequent activation of the HPA axis, resulting in increased FKBP51 production, can lead to GR resistance, diminished negative feedback on the HPA axis, and prolonged stress hormone activation ^35^. Accordingly, *FKBP5* gene variants, that can influence GR sensitivity to cortisol ^36^, were associated with depression and anxiety disorders ^36–41^, as well as with substance use disorders ^42–50^.

Although the effect of stress and the HPA axis on memory processes has been thoroughly investigated, most of this research has focused on fear memories ^51,52^, with only a few studies addressing drug-related memories ^53–55^. Here, we sought to explore the involvement of the HPA axis, and specifically of FKBP51, in the modulation of alcohol-related memory reconsolidation. To this end, we first investigated the effect of FKBP51 inhibition on the reconsolidation of contextual alcohol memories. Then, we characterized the effect of chronic alcohol exposure on *Fkbp5* responsiveness to alcohol-related cues and acute alcohol exposure. Finally, we tested the effect of alcohol memory retrieval on corticosterone secretion and investigated the causal role of the HPA axis in memory reconsolidation using behavioral and pharmacological manipulations.

## 2. Materials and methods

### 2.1. Animals

Male and female C57BL/6J mice and Wistar rats (18-25 gr and 160-240 gr at the beginning of the experiments, respectively) were bred at Tel-Aviv University animal facility, and housed under a 12-h light-dark cycle (lights on at 7 a.m.) with food and water available ad libitum. Animals were individually housed, except for conditioned-place preference (CPP) experiments, in which mice were housed 2-4 per cage. Animals were weighed once a week to control for weight loss, except for CPP experiments, in which animals were weighed twice a week. All experimental protocols were approved by and conformed to the guidelines of the Institutional Animal Care and Use Committee of Tel Aviv University, and the NIH guidelines (animal welfare assurance number A5010-01). All efforts were made to minimize the number of animals used.

### 2.2. Drugs and Reagents

Ethanol absolute (0005250502000) (BioLab, Jerusalem, Israel) was diluted to a 20% (v/v) solution with tap water for voluntary drinking, or with saline solution (0.9 M NaCl) for systemic injections. Isoflurane was obtained from Piramal Critical Care (Bethlehem, PA, USA). The FKBP51 inhibitor, SAFit2 ^56^ (supplied by the lab of Felix Hausch, Technical University Darmstadt, Germany), was solubilized in 4% DMSO, 2% Tween-80, and saline. The glucocorticoid receptor antagonist, mifepristone (459980010) (Thermo Scientific Chemicals, Waltham, MA, USA), was solubilized in 100% DMSO. Corticosterone (27840) (Merck, Rehovot, Israel) was solubilized in 5% DMSO and saline. DetectX Corticosterone Enzyme Immunoassay Kit (K014) was supplied by Arbor Assays (Ann Arbor, MI, USA). Fast SYBR Green Master Mix (4385610), TRIzol reagent (15596026), and RevertAid kit (K1691) were supplied by Thermo-Fisher Scientific (Waltham, MA, USA). DNA oligonucleotides (RT-qPCR primers) were obtained from Merck (Rehovot, Israel).

### 2.3. Behavioral Procedures

#### 2.3.1. Alcohol-Conditioned Place Preference (CPP)

##### Apparatus

Place conditioning experiments were performed in open ceiling Plexiglas boxes (30 x 20 x 20 cm) divided into two equal-sized compartments by a white sliding door. The compartments differed in wall pattern (horizontal vs. vertical black and white stripes, 1 cm wide) and the texture of the silicone floor surface (bulging circles vs. bulging stripes). The horizontal stripes pattern of the walls was coupled with the circles floor texture, and the vertical stripes pattern of the walls was coupled with the stripes floor texture. Each Plexiglas box was placed in a sound-attenuating chamber equipped with a LED light stripe on the walls and a ceiling camera that registered mice behavior. Each CPP chamber was assigned to a single sex. Data were recorded and analyzed with Ethovision XT 11.5 (Noldus, Wageningen, Netherlands).

##### CPP procedure

The CPP procedure was performed as previously described ^25^ and began after 3-5 days of habituation to i.p. saline injections.

##### Baseline test (day 1)

on the first day, the sliding door between the two compartments of the apparatus was elevated, and mice were allowed to freely explore the apparatus for 30 min. The time spent in each compartment was recorded, and mice who spent >70% of the time in either of the compartments were excluded from the experiment (three mice in Experiment 1, one mouse in Experiment 6, one mouse in Experiment 7, three mice in Experiment 8). This allowed an unbiased design, in which the pre-conditioning preference for each of the compartments, as indicated by the group average, is equal ^57^.

##### Place conditioning (days 2-9)

over eight consecutive days mice were injected with saline (days 2, 4, 6, and 8) or with alcohol (1.8 gr/kg, 20% v/v, i.p.) (days 3, 5, 7 and 9). Immediately after the injection, the mice were placed for five min in one of the CPP apparatus compartments, with the sliding door closed. Each compartment was paired either with saline or with alcohol in a counterbalanced manner. On the days of the alcohol injections the doors of the chambers in which the CPP boxes were placed were sprayed with 70% alcohol solution, to create an odor cue.

##### Pre-retrieval place-preference test (day 10)

twenty-four hours after the last conditioning session, mice were subjected to a place-preference test, during which they were placed in the CPP apparatus, with the sliding door elevated, and were allowed to freely explore the apparatus for 15 min. The alcohol place-preference score was indexed as the percentage of time spent in the alcohol-paired compartment (time spent in the alcohol-paired compartment × 100 / total test time). More time spent in the alcohol-paired compartment indicates higher alcohol seeking. Mice that failed to show >5% increase in their preference for the alcohol-paired compartment, compared with the Baseline test, were excluded from the experiment (20 mice in Experiment 1, 16 mice in Experiment 6, 16 mice in Experiment 7, nine mice in Experiment 8, 12 mice in Experiment 9).

##### Memory retrieval (day 11)

prior to this stage, the mice were assigned to the different experimental conditions (matched to the CPP scores of the baseline and the pre-retrieval tests). In order to retrieve the alcohol memories, mice were placed for three min in the compartment that was paired with alcohol during the conditioning phase, when the sliding door of the CPP apparatus is closed, and in the presence of alcohol odor. In experiments with pharmacological or behavioral manipulations, all the mice underwent the retrieval process and were re-exposed to the alcohol-paired compartment. In experiments that involved post-retrieval molecular measurements, mice were either exposed to the alcohol-paired compartment (Retrieval group) or were briefly handled (No Retrieval group).

##### Post-retrieval place preference tests (extinction) (days 12-15)

in experiments that involved pharmacological or behavioral manipulations, changes in the mice preference to the alcohol-paired compartments were monitored in four post-retrieval (extinction) tests, 24 hours apart, which were identical to the pre-retrieval place preference test.

#### 2.3.2. Restraint stress

The restraint stress procedure was conducted as previously described ^58^. Briefly, mice were placed for 30 min in a 50 ml falcon tube with ventilation holes. The tube allows enough space for back- and forth-movement, but not for turning around. Control mice were left untouched in their home cages.

#### 2.3.3. Intermittent access to 20% alcohol in 2-bottle choice (IA2BC)

After one week of habituation to individual cages, mice or rats were trained to consume 20% alcohol solution in the IA2BC as previously described ^59–61^. Animals were given three 24-hour sessions of free access to 2-bottle choice per week (tap water and 20% alcohol v/v) on Sundays, Tuesdays, and Thursdays. Alcohol drinking sessions were followed by a 24-or 48-hour deprivation period, in which the animals received only water, producing repeated cycles of intoxication and withdrawal. The position (left or right) of the alcohol and water bottles was alternated between the sessions to control for side preference. Alcohol exposure in this protocol was shown to produce high levels of alcohol intake and blood alcohol concentration, especially in the first hours of drinking ^59–63^. Water and alcohol bottles were weighed before and after each alcohol-drinking session, and consumption levels were normalized to body weight. Training lasted 7-8 weeks for mice, and 8-10 weeks for rats.

##### Memory retrieval after IA2BC

following 8-10 weeks of training, rats were subjected to 10 days of abstinence. On the 11^th^ day of abstinence, alcohol memory retrieval was conducted at the rats’ home cages with 10 min exposure to an empty bottle of alcohol with its tip covered with a drop of alcohol, serving as an odor-taste cue, as was previously described ^20,64^. Control rats were left untouched in their home cages.

### 2.4. Blood collection and plasmatic corticosterone assessment

Mice were anesthetized continuously with Isoflurane and blood was collected by cardiac puncture using EDTA-coated syringes (1.8 mg of EDTA dissolved in 40 ul of PBS). Plasma was separated by centrifugation at 1000 G for 20 min at 4°C and was later stored at -80°C. Plasma corticosterone was measured using the DetectX Corticosterone Enzyme Immunoassay Kit, according to the manufacturer’s instructions, with all samples ran in duplicates. Five ul from each sample were processed for the analysis, to reach a final dilution of 1:80. The optical density readings were analyzed using the online tool MyAssays, and the calculated corticosterone concentration values were adjusted according to the *volume of collected blood / volume of EDTA solution* ratio. The averaged intra-assay coefficient of variance (CV%) was 2%.

### 2.5. Quantitative reverse transcriptase polymerase chain reaction (qRT-PCR)

Following brain dissection, tissue samples were immediately snap-frozen in liquid nitrogen and stored at −80°C until use. Frozen tissues were mechanically homogenized in TRIzol reagent, and total RNA was isolated from each sample according to the manufacturer’s instructions. mRNA was reverse transcribed to cDNA using the Reverse Transcription System and RevertAid kit. Plates (96 wells) were prepared for SYBR Green cDNA analysis using Fast SYBR Master Mix. Samples were analyzed in duplicate with a Real-Time PCR System (StepOnePlus, Applied Biosystems, Foster City, CA, USA). The following reaction primers sequences were used: *Fkbp5* (mice) forward, 5′- ATTTGATTGCCGAGATGTG -3′; reverse, 5′- TCTTCACCAGGGCTTTGTC -3′; *Fkbp5* (rats) forward, 5′- TCCTTGTCAGCGTGTGTACC -3′; reverse, 5′- GAGCGAGGTATCTGCCTGTC -3′; *Gapdh* (mice) forward, 5′- CCAGAACATCATCCCTGC -3′; reverse, 5′- GGAAGGCCATGCCAGTGAGC-3′; *Gapdh* (rats) forward, 5′- GCAAGAGAGAGGCCCTCAG -3′; reverse, 5′- TGTGAGGGAGATGCTCAGTG -3′. Thermal cycling was initiated with incubation at 95°C for 20 sec, followed by 40 cycles of PCR with the following conditions: Heating at 95°C for 3 sec and then 30 sec at 60°C. Relative quantification was calculated using the ΔΔCt method. The expression of target genes was normalized to that of *Gapdh* as an internal control gene ^61^. Change in the mRNA expression of the experimental groups was calculated as the percentage of the control group.

### 2.6. Experimental design and statistical analysis

The allocation of mice and rats to the experimental groups was done based on their average CPP scores (in the CPP experiments) or alcohol drinking (in the IA2BC experiments), to create groups with approximately equal place preference or alcohol drinking levels. Sex was distributed approximately equally across the experiments and was initially analyzed as a factor. Aside from a single experiment (Experiment 1), the analyses in all other experiments did not yield an interaction between sex and other factors (p’s > 0.05). Therefore, data were collapsed across this factor.

#### 2.6.1. Experiment 1 – Effects of FKBP51 inhibition on alcohol memory reconsolidation

Mice were trained in the CPP paradigm as described above. The FKBP51 inhibitor SAFit2 (20 mg/kg, i.p., injection volume 10 ml/kg) or vehicle was injected 16 hours before the alcohol memory retrieval session. This interval was chosen based on previous studies that tested the anxiolytic properties of SAFit2 and reported an anxiolytic effect of SAFit2 16 hours, but not one hour, after the treatment ^56,65^. The mice’s preference for the alcohol-paired compartment was monitored over the four following days in daily place preference tests. Expression of alcohol-CPP following conditioning was analyzed by a paired t-test (baseline test vs. pre-retrieval place-preference test). Place preference in the post-retrieval tests was analyzed by a 3-way mixed-model ANOVA, with between-subjects factors of Treatment (SAFit2, vehicle) and Sex (males, females), and a repeated measures factor of Post-retrieval test (post-retrieval test 1-4). Significant ANOVA was followed by contrast analysis.

#### 2.6.2. Experiments 2+3 – Effects of alcohol memory retrieval on Fkbp5 mRNA expression in a 2-bottle choice procedure

The paradigm of alcohol memory retrieval in the home cage, after a period of voluntary alcohol drinking, was previously established in rats’ experiments ^20^. Therefore, in this experiment, we used the rat model. Rats were trained to voluntarily consume alcohol in the IA2BC protocol, as described above. Following 8-10 weeks of drinking, rats were subjected to 10 days of abstinence. On the 11^th^ day of the abstinence period, the rats’ alcohol memory was retrieved, using an odor-taste cue, as described above. Control rats did not undergo the memory-retrieval procedure and had access to water bottles. Ten min or 30 min (Experiment 2), and 60 min (Experiment 3) after the memory retrieval manipulation, the rats were euthanized, and brain tissues were processed for qRT-PCR analysis. Three regions were examined: the mPFC, in which we previously found alterations in *Fkbp5* expression after alcohol memory retrieval in the CPP paradigm ^25^, the amygdala, which was shown to have a critical role in the modulatory effect of the HPA axis on reconsolidation of emotional memories ^52^, and drug memories in particular ^54,55^, and the nucleus accumbens (NAc), which was shown to be critical in cue- and context-induced reinstatement to alcohol-seeking behavior ^66–68^. The mRNA expression of *Fkbp5* for each brain region was normalized to *Gapdh*, and is expressed as the percentage of the no-retrieval control group’s expression. In Experiment 2, *Fkbp5* expression data from each brain region were analyzed by one-way ANOVA, with a between factor of Group (no-retrieval, retrieval-10 min, retrieval-30 min). mRNA expression data in Experiment 3 were analyzed by an unpaired t-test (no retrieval vs. retrieval 60 min).

#### 2.6.3. Experiment 4 – Effects of alcohol consumption on Fkbp5 mRNA expression

The IA2BC drinking protocol produces higher levels of alcohol consumption and preference in mice, compared with rats ^69^. Therefore, the use of mice for assessing neuroadaptations that arise due to prolonged alcohol consumption is preferable. Here, mice were trained to voluntarily consume alcohol in the IA2BC protocol, as described above. Control mice were exposed to water only. Following eight weeks of drinking, the mice were euthanized at three different time points: four hours into the last drinking session (9 p.m.) (binge-like drinking), at the end of the drinking session, and after 24 hours of withdrawal. The tissue collection from the control mice was divided between the three time points, and as the expression data in the different control intervals did not differ, data were collapsed across intervals in vehicle controls. Brain tissues were processed for qRT-PCR analysis. The *Fkbp5* mRNA expression for each brain region was normalized to *Gapdh* and is expressed as the percentage of the water group’s expression. *Fkbp5* expression data from each brain region were analyzed by one-way ANOVA, with a between factor of Group (water, alcohol 4 hours, alcohol 24 hours, alcohol withdrawal). Significant ANOVA was followed by Dunnett post-hoc test. The mice’s averaged alcohol consumption (over the last five sessions) was correlated with the *Fkbp5* expression in the amygdala, where a change in the expression was observed in the 4 hours time point, using Pearson’s correlation.

#### 2.6.4. Experiment 5 – Effects of alcohol injection, with or without a history of long-term alcohol drinking, on Fkbp5 mRNA expression

Mice were trained to voluntary consume alcohol in the IA2BC protocol, as described above. Control mice were exposed to water only. Following eight weeks of training, mice were injected with alcohol (1.8 gr/kg, 20% v/v, i.p.) or with saline, to form four experimental groups: water drinking history + saline challenge, water drinking history + alcohol challenge, alcohol drinking history + saline challenge, and alcohol drinking history + alcohol challenge. Before the challenge test day, the mice received three intermittent sessions of injection handling. Brain tissues were collected two hours after the alcohol/saline challenge injection and were processed for qRT-PCR analysis. The mRNA expression for each brain region was normalized to *Gapdh* and is expressed as the percentage of the water+saline control group’s expression. mRNA expression data from each brain region were analyzed by two-way ANOVA, with Challenge (alcohol, saline) and Drinking history (alcohol, water) as between factors.

#### 2.6.5. Experiment 6 – Effects of alcohol memory retrieval on plasma corticosterone levels

Mice were trained in the CPP paradigm as described above. On the 11^th^ day of the protocol, the alcohol memory was retrieved as described above. Control mice did not undergo the memory-retrieval procedure and instead were briefly handled. Ten min or 30 min after alcohol-memory retrieval, blood samples were collected and processed for plasmatic corticosterone assessment. Expression of alcohol-CPP following conditioning was analyzed by a paired t-test (baseline test vs. pre-retrieval place-preference test). Corticosterone levels were analyzed by a two-way ANOVA test with between-factors of Group (retrieval, no-retrieval) and Time after retrieval (10 min, 30 min).

#### 2.6.6. Experiment 7 – Effects of restraint stress on alcohol memory reconsolidation

Mice were trained in the CPP paradigm, and 10 min after the alcohol-memory retrieval session they were subjected to 30 min of restraint stress as described above. Importantly, the restraint-stress procedure was previously shown to induce increased secretion of the adrenocorticotropic hormone and corticosterone ^70^. The mice’s preference for the alcohol-paired compartment was monitored over the four following days in daily place preference tests. The expression of alcohol-CPP following conditioning was analyzed by a paired t-test (baseline test vs. pre-retrieval place-preference test). Place preference in the post-retrieval tests was analyzed by a mixed-model ANOVA, with a between-subjects factor of Group (stress, no stress) and a repeated measures factor of Post-retrieval test (post-retrieval test 1-4).

#### 2.6.7. Experiment 8 – Effects of corticosterone on alcohol memory reconsolidation

Mice were trained in the CPP paradigm, and immediately after the alcohol-memory retrieval session, they were injected with corticosterone (1 or 10 mg/kg, i.p., injection volume 10 ml/kg), or with vehicle solution. The mice’s preference for the alcohol-paired compartment was monitored over the four following days in daily place preference tests. Expression of alcohol-CPP following conditioning was analyzed by a paired t-test (baseline test vs. pre-retrieval place-preference test). Place preference in the post-retrieval tests was analyzed by a mixed-model ANOVA, with a between-subjects factor of Treatment (corticosterone, vehicle) and a repeated measures factor of Post-retrieval test (post-retrieval test 1-4).

#### 2.6.8. Experiment 9 – Effects of glucocorticoid receptor inhibition on alcohol memory reconsolidation

Mice were trained in the CPP paradigm, and immediately after the alcohol-memory retrieval session, they were injected with the glucocorticoid receptor antagonist, mifepristone (10 or 60 mg/kg, i.p., injection volume 2 ml/kg), or with vehicle solution. The mice’s preference for the alcohol-paired compartment was monitored over the four following days. Expression of alcohol-CPP following conditioning was analyzed by a paired t-test (baseline test vs. pre-retrieval place-preference test). Place preference in the post-retrieval tests was analyzed by a mixed-model ANOVA, with a between-subjects factor of Treatment (mifepristone, vehicle) and a repeated measures factor of Post-retrieval test (post-retrieval test 1-4).

## 3. Results

### FKBP51 inhibition before alcohol-memory retrieval enhances memory stability in female mice

We recently showed that *Fkbp5* mRNA expression is increased following the retrieval of alcohol memories in the alcohol-CPP paradigm ^25^. Here, we tested whether FKBP51 plays a causal role in alcohol-memory reconsolidation.

Mice were trained in the alcohol-CPP paradigm to prefer the alcohol-associated compartment (see Methods). Upon the formation of the alcohol-associated memories, they were retrieved by briefly re-exposing the mice to the alcohol-paired context. The FKBP51 inhibitor SAFit2 or vehicle were injected 16 hours before the memory-retrieval session (20 mg/kg, i.p., dose based on Hartmann et al. ^65^), as SAFit2 was previously shown to exert its anxiolytic effects, which were also reflected in reduced alcohol consumption of rats with a history of stress exposure ^71^, 16 hours after administration ^56,65^. Alcohol-seeking behavior was then monitored in four post-retrieval tests under extinction conditions (no alcohol given), 24 hours apart (Figure 1A) (see Experiment 1 in Methods – Experimental Design).

**Figure 1.**
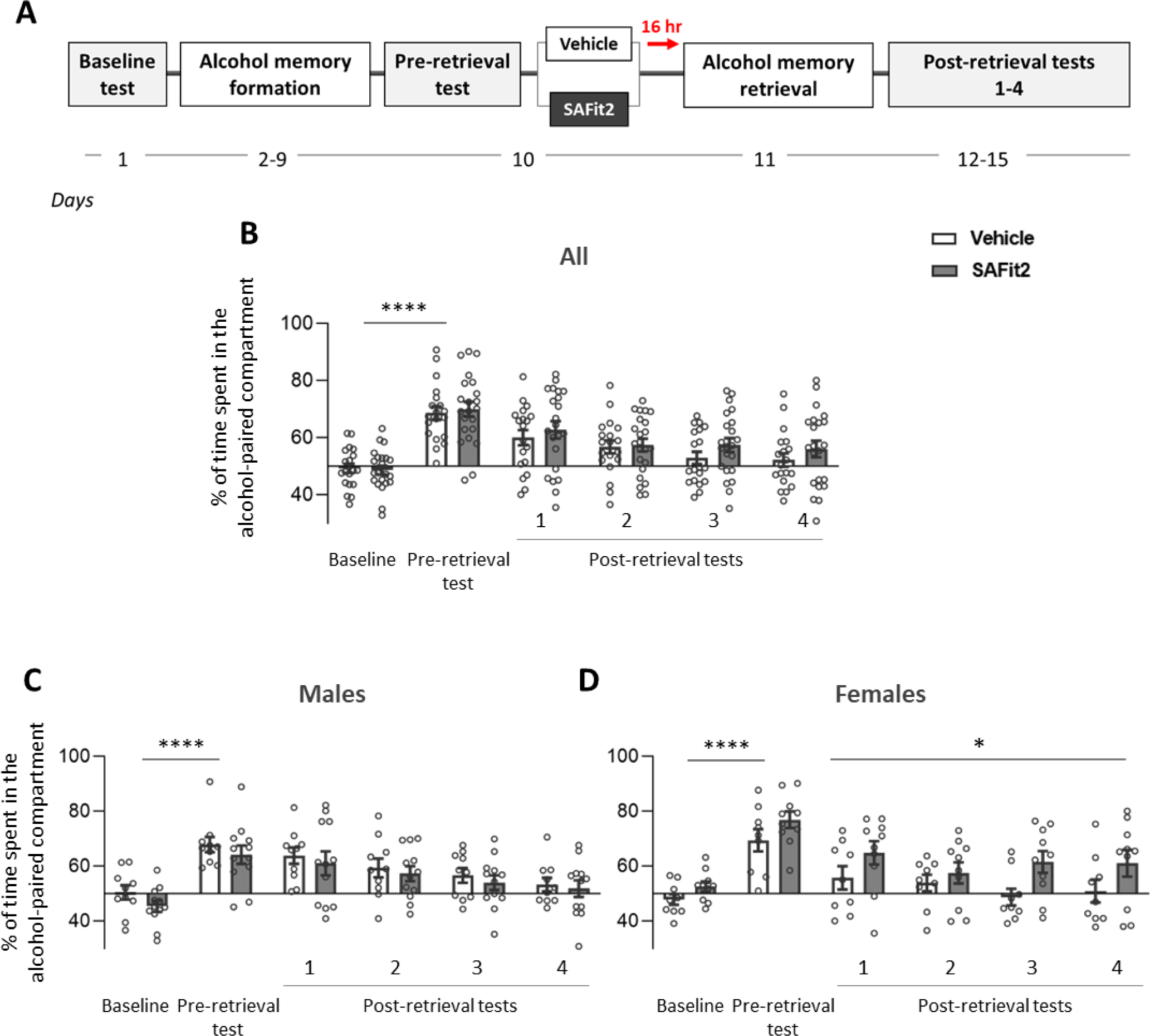
Treatment with the FKBP51 inhibitor SAFit2 before alcohol-memory retrieval increases the stability of the alcohol memory in females, but not in males. **A.** Experimental design and timeline. Mice were trained in an alcohol CPP procedure and were then treated with SAFit2 (20 mg/kg) or vehicle 16 hours before the retrieval of the alcohol memories. The mice’s preference for the alcohol-paired compartment was monitored in four subsequential tests, 24 hours apart. B-D. Bar graphs indicate the percent of time spent in the alcohol-paired compartment of all the mice (B), males (C), and females (D). Bar graphs represent the mean±S.E.M. Males: n=10-12; Females: n=9-10. ****p < 0.0001 (Baseline test vs. Pre-retrieval test); * p<0.05 (Vehicle vs. SAFit2).

As expected, the mice displayed a higher preference for the alcohol-paired compartment after the conditioning phase, compared to their baseline preference (Figure 1; paired t-test: All mice: t(40)=11.99; Males: t(21)=7.89; Females: t(18)=8.09; p’s<0.0001). We found that the repeated tests led to an extinction effect (reduction in the preference of the alcohol compartment) in the vehicle-treated control group, as expected. However, SAFit2-treated mice showed a weaker, partial extinction effect (Figure 1B). When dividing the mice by sex, we found that the extinction-attenuating effects of SAFit2 were observed in females, but not in males (Figures 1C-D) (mixed model ANOVA for All mice: A main effect of Test [F(3,111)=7.32, p<0.001], marginally significant Sex X Treatment interaction effect [F(1,37)=3.6, p=0.066]; but no main effects of Treatment [F(1,37)=1.29, p=0.264] or Sex [F(1,37)=0.02, p=0.901] and no Test X Treatment [F(3,111)=0.65, p=0.586], Test X Sex [F(3,111)=1.40, p=0.248], and Test X Treatment X Sex interaction [F(3,111)=0.75, p=0.526]; Contrast analysis: a significant difference between the vehicle and SAFit2 groups in the post-retrieval tests in female [F(1,37)=4.29, p=0.045], but not in male mice [F(1,37)=0.31, p=0.580]).

These results suggest that in females, inhibition of FKBP51 adjacent to the retrieval of alcohol memory prevents the extinction of alcohol CPP and enhances memory stability during its reconsolidation. This effect results in prolonged alcohol-seeking behavior.

### Fkbp5 mRNA expression is not affected by the retrieval of alcohol memories in a long-term alcohol self-administration model in rats

The CPP paradigm is widely used to investigate contextual memory processes and drug/alcohol reward, but alcohol exposure in this paradigm is moderate and involuntary, and does not model drug abuse. Therefore, we next investigated the effect of alcohol memory retrieval on *Fkbp5* expression in a behavioral paradigm more relevant to alcohol abuse and addiction, namely, long-term voluntary alcohol self-administration in the IA2BC procedure in rats, as we previously described ^20^.

We trained rats to drink alcohol in the IA2BC paradigm (see Methods) for 8-10 weeks. The average alcohol intake during the last five drinking sessions was 5.17 gr/kg (SD=2.08) in Experiment 2, and 4.08 gr/kg (SD=1.72) in Experiment 3. Rats were then subjected to 10 days of abstinence, and on the 11^th^ day, alcohol memories were retrieved with an odor-taste cue ^20,64^ (see Methods). mPFC, amygdala, and NAc tissues were collected 10 and 30 min (Experiment 2) or 60 min (Experiment 3) after alcohol-memory retrieval (Figure 2A).

**Figure 2.**
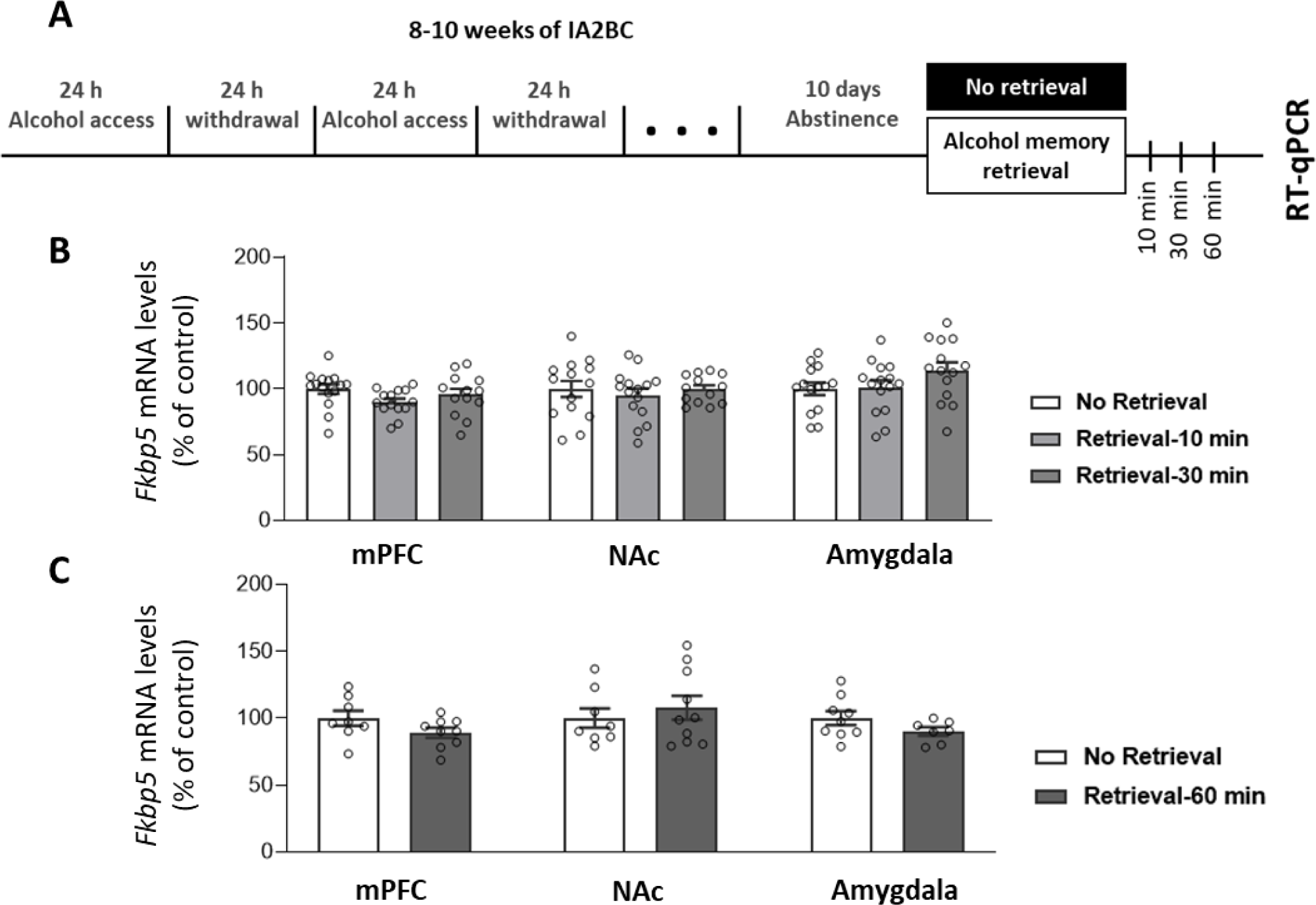
Alcohol-memory retrieval after voluntary alcohol consumption has no effect on *Fkbp5* mRNA expression in rats. **A.** Experimental design and timeline. Rats consumed alcohol in the intermittent access to 20% alcohol 2-bottle choice (IA2BC) paradigm for 8-10 weeks, followed by 10 days of abstinence. After the abstinence period, alcohol memory was retrieved using an odor-taste cue. In one experiment, tissues were collected 10 and 30 min after the retrieval, and in the second experiment, tissues were collected 60 min after the retrieval. *Fkbp5* mRNA levels were determined by RT-qPCR, and normalized to *Gapdh.* **B-C.** *Fkbp5* expression in the mPFC, the NAc, and the amygdala, 10, 30 (B) or 60 (C) min after the retrieval. Bar graphs represent the mean±S.E.M. of the percent of change from the control group. 10/30 min experiment: n=7-10; 60 min experiment: n=13-15.

As shown in Figures 2B-C, we observed no changes in the mRNA levels of *Fkbp5* in any of the brain regions and time points (10/30 min experiment: one-way ANOVA: mPFC [F(2,39)=2.04, p=0.143]; NAc [F(2,39)=0.29, p=0.746]; Amygdala [F(2,40)=1.94, p=0.156]; 60 min experiment: unpaired t test: mPFC [t(15)=1.64, p=0.122]; NAc [t(16)=0.66, p=0.518]; Amygdala [t(14)=1.52, p=0.151]).

These results indicate that while retrieval of alcohol memory using contextual cues leads to increased expression of *Fkbp5*, retrieval using an odor-taste cue, and after a period of voluntary alcohol consumption, has no effect on *Fkbp5* expression.

### Chronic-voluntary alcohol consumption does not affect Fkbp5 mRNA expression in mice

Our findings that alcohol memory retrieval increases *Fkbp5* expression in the CPP paradigm ^25^, but not in the IA2BC self-administration paradigm, raise the possibility that the prolonged voluntary exposure to alcohol during the IA2BC procedure has possibly made *Fkbp5* less responsive to alcohol and cues associated with it. To test this possibility, we first tested whether alcohol exposure in this manner would affect *Fkbp5* mRNA expression in the mPFC, NAc, and amygdala.

Mice were trained to consume alcohol in the IA2BC procedure as described above, with control mice consuming water only (see Methods). Tissues were collected after seven weeks of alcohol drinking, at three different time points: after the first four hours of the last drinking session (binge-like drinking), at the end of the 24-hour drinking session, and after 24 hours of withdrawal (Figure 3A) (see Experiment 4 in Methods). As depicted in Figure 3B, the average alcohol intake during the last five drinking sessions was 13.25 gr/kg (SD=5.02).

**Figure 3.**
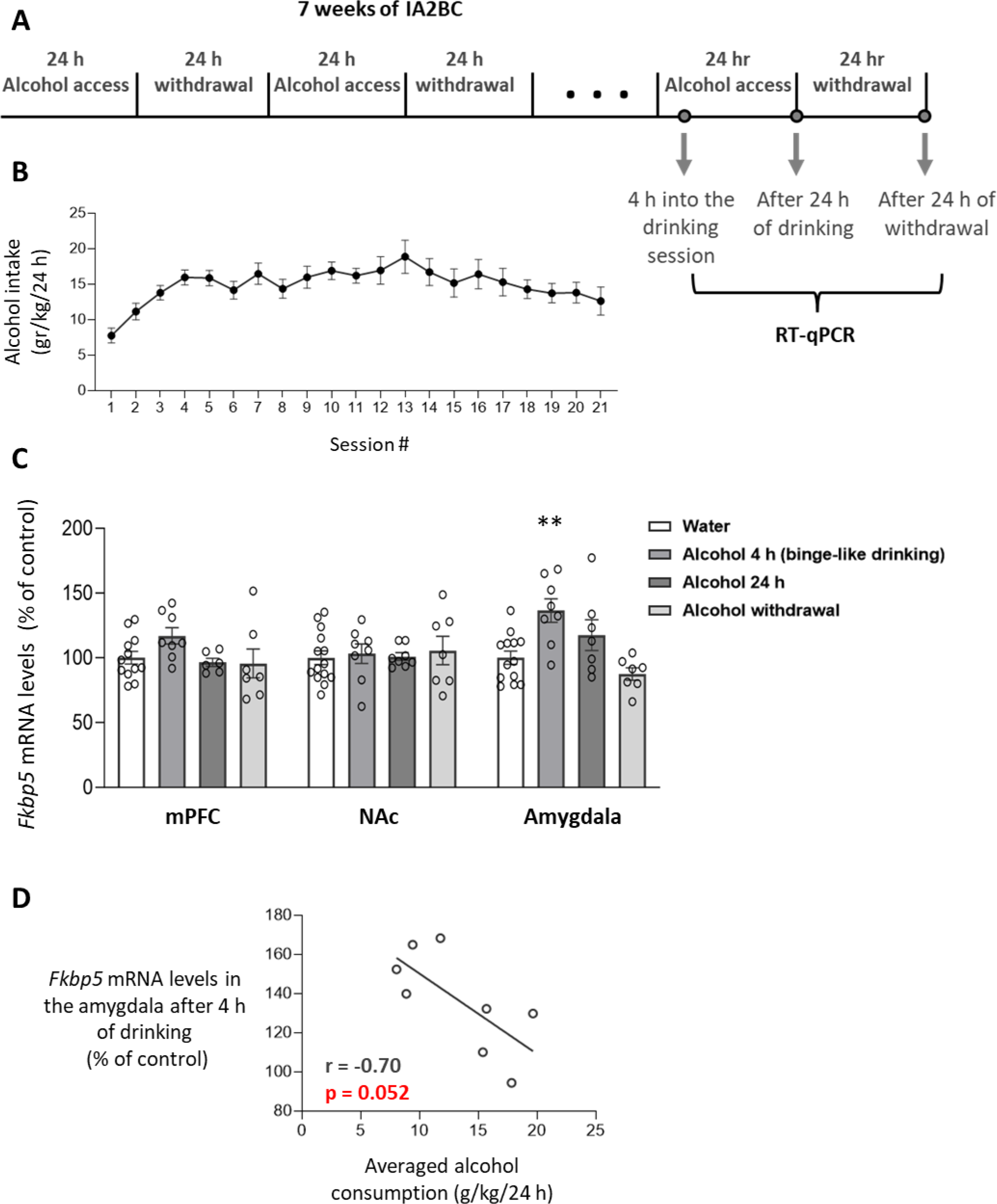
Voluntary alcohol consumption increases *Fkbp5* expression in the amygdala of mice. **A.** Experimental design and timeline. Mice consumed alcohol in the intermittent access to 20% alcohol 2-bottle choice (IA2BC) paradigm for seven weeks (21 sessions), and alcohol consumption was measured daily. Control mice received access to water only. Tissues were collected four hours into the last drinking session (binge-like drinking), after 24 hours of drinking, and after 24 hours of withdrawal. *Fkbp5* mRNA levels were determined by RT-qPCR, and normalized to *Gapdh.* **B.** Alcohol intake throughout the 21 IA2BC drinking sessions (mean±S.E.M)**. C.** *Fkbp5* expression in the mPFC, the NAc, and the amygdala. Bar graphs represent the mean±S.E.M. of the percent of change from the control group. **D.** Correlation between the average alcohol consumption during the last five drinking sessions and *Fkbp5* expression in the amygdala after four hours of alcohol drinking (binge-like drinking). Control groups: n=12-14, alcohol groups: n=6-8. ** p<0.01 compared with water-drinking control.

As shown in Figure 3C, *Fkbp5* levels in the amygdala were increased after binge-like alcohol drinking (4 h time point), compared with water-drinking controls, with a similar trend for the 24 h drinking group in the amygdala (one-way ANOVA: F(3,31)=7.21, p<0.001. post hoc, binge vs. water: p=0.003). No effects were found in the mPFC and the NAc (one-way ANOVA: mPFC [F(3,29)=2.02, p=0.133]; NAc [F(3,33)=0.13, p=0.942]).

Interestingly, the average alcohol consumption during the last five drinking sessions was negatively correlated with the *Fkbp5* mRNA expression level in the amygdala, with mice that displayed higher alcohol consumption showing lower *Fkbp5* expression at the binge-like drinking point (Pearson’s r = -0.70, p=0.052) (Figure 3D). In summary, our results show that while binge-like drinking increases *Fkbp5* expression in the amygdala, the global consumption is negatively associated with the expression of the gene, and alcohol drinking does not affect the expression in the mPFC or the NAc.

### Acute injection of alcohol leads to increased expression of Fkbp5 mRNA in mice, regardless of alcohol-drinking history

The negative correlation that was observed between the mice’s *Fkbp5* expression in the amygdala at the binge-like drinking time point and their averaged alcohol consumption, may suggest that the higher the alcohol consumption was, the smaller the increase in *Fkbp5* is. We, therefore, hypothesized that a prolonged period of alcohol drinking has turned *Fkbp5* transcription less responsive to alcohol. Therefore, we next examined whether a history of alcohol consumption affects *Fkbp5’s* responsiveness to alcohol exposure.

Mice consumed alcohol as described above, with controls consuming only water. The average alcohol intake during the last five drinking sessions was 13.78 gr/kg (SD=6.05). After eight weeks of heavy alcohol consumption, following 24 hours of withdrawal, mice were injected with an alcohol challenge injection (1.8 gr/kg, i.p.) or saline. Brain tissues were collected two hours after the alcohol/saline challenge (Figure 4A) (see Experiment 5 in Methods). *Fkbp5* mRNA levels were determined using qRT-PCR.

**Figure 4.**
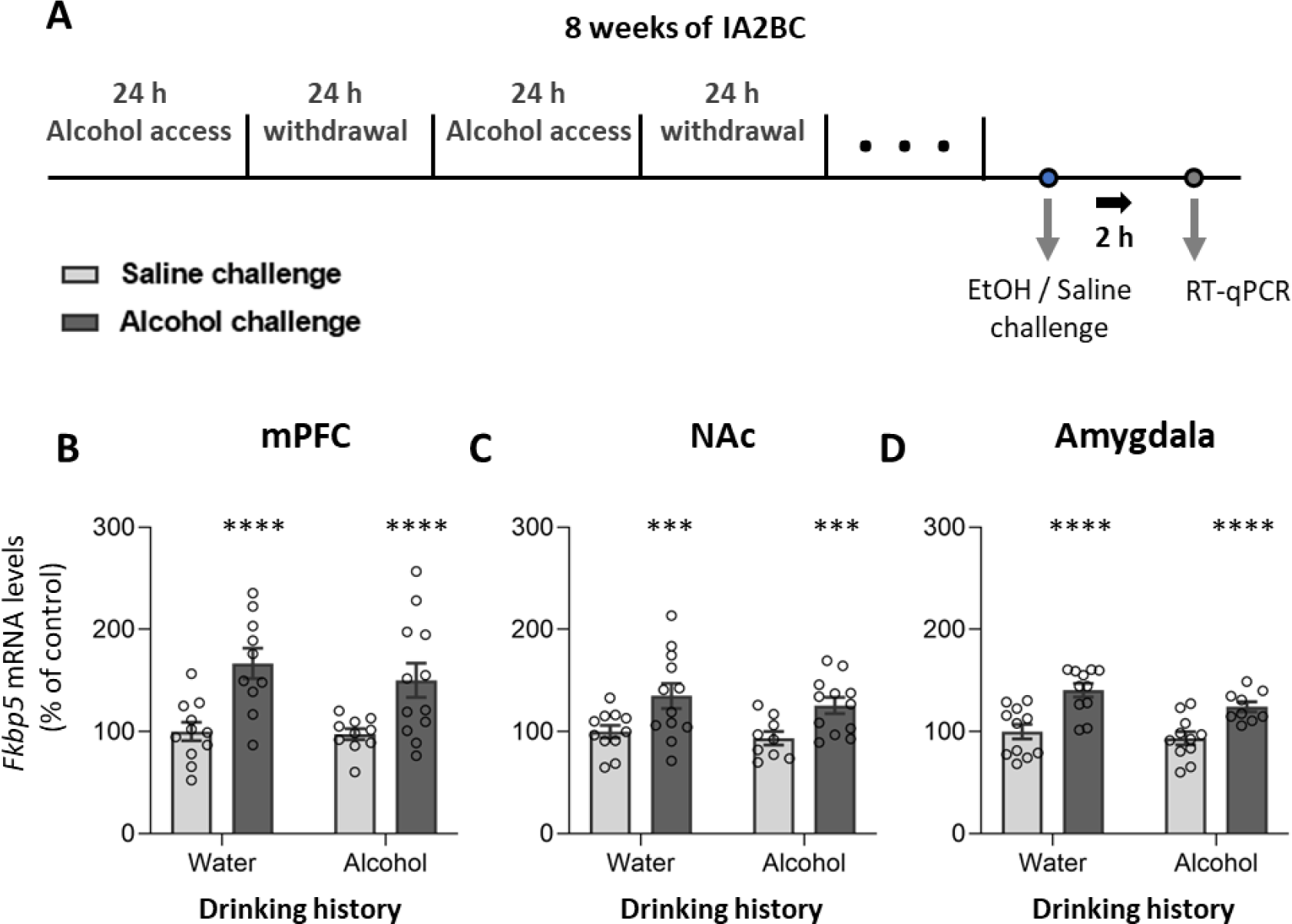
Alcohol injection increases *Fkbp5* expression regardless of alcohol drinking history. **A.** Experimental design and timeline. Mice consumed alcohol in the intermittent access to 20% alcohol 2-bottle choice paradigm. After eight weeks of drinking, mice were challenged with an injection of saline or alcohol (1.8 mg/kg, i.p.). The tissues were collected two hours after the injection. *Fkbp5* mRNA levels were determined by RT-qPCR, and normalized to *Gapdh.* **B-D.** *Fkbp5* expression in the mPFC (B), NAc (C), and amygdala (D). Bar graphs represent the mean±S.E.M. of the percent of change from the control group. n=9-12. *** p<0.001, **** p<0.0001 (alcohol challenge vs. Saline challenge).

As shown in Figure 4B-D, the alcohol challenge injection led to increased expression of *Fkbp5* in the mPFC, the NAc and the amygdala, regardless of the alcohol drinking history (Two-way ANOVA: a main effect for Challenge [mPFC [F(1,39)=21.67, p<0.0001]; NAc [F(1,40)=13.72, p<0.001]; amygdala [F(1,38)=29.58, p<0.0001]], but no main effect for Drinking history [mPFC [F(1,39)=0.55, p=0.462]; NAc [F(1,40)=0.76, p=0.389]; amygdala [F(1,38)=2.99, p=0.092]], and no Challenge X Drinking history interaction [mPFC [F(1,39)=0.31, p>0.583]; NAc [F(1,40)=0.02, p=0.876]; amygdala [F(1,38)=0.53, p=0.471]]). These findings suggest that *Fkbp5* responsiveness to alcohol exposure is not affected by a history of chronic alcohol consumption.

### Retrieval of contextual alcohol memory leads to elevated corticosterone levels in the plasma

Corticosterone is a positive regulator of *Fkbp5* expression ^27,30^. Since we previously identified increased expression of *Fkbp5* mRNA after alcohol-memory retrieval ^25^, we sought to characterize changes in the plasma levels of corticosterone following this procedure. Mice were trained in the CPP procedure as described above. Upon the formation of the alcohol-associated memories, they were retrieved by briefly re-exposing the mice to the alcohol-paired context, with no-retrieval controls being handled. We assessed the plasma levels of corticosterone 10 or 30 min after the alcohol memory retrieval (Figure 5A) (see Experiment 6 in Methods).

**Figure 5.**
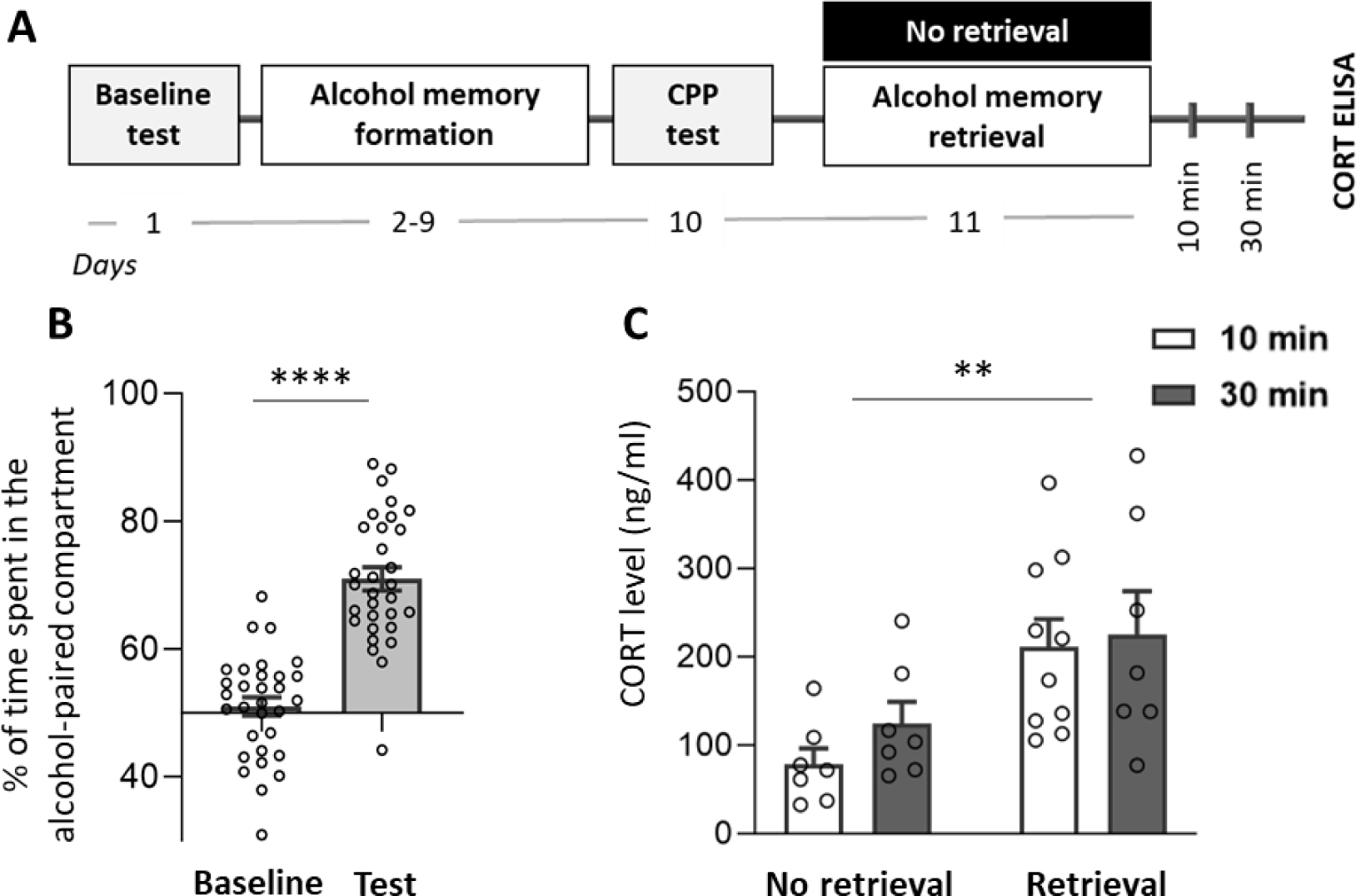
Alcohol memory retrieval increases corticosterone secretion. **A.** Experimental design and timeline. Mice underwent place conditioning to alcohol in the CPP paradigm. Memory retrieval was done using a 3-minute exposure to the context that was previously associated with alcohol. Control mice were brought to the procedure room and underwent short handling but were not re-exposed to the alcohol-paired context. Blood was withdrawn 10 and 30 min after the alcohol memory retrieval. Corticosterone levels in the plasma were determined using ELISA. **B.** Bar graphs indicate the mean of the percent of time spent in the alcohol-paired compartment ± SEM. **C.** Bar graphs represent mean±S.E.M. of corticosterone concentration n=7-10. **** p<0.0001 (Baseline vs. Test), ** p<0.01 (No retrieval vs. Retrieval).

As expected, after the conditioning phase, the mice displayed a higher preference for the alcohol-paired compartment compared to their preference in the baseline session (Figure 5B; paired t-test: t(30)=11.59; p<0.0001). As depicted in Figure 5C, the retrieval group showed higher levels of plasma corticosterone, compared to no-retrieval controls (two-way ANOVA: a main effect of Group [F(1,27)=12.40, p=0.002], but no main effect for Time after retrieval [F(1,27)=0.81, p=0.376], and no Group X Time after retrieval interaction effect [F(1,27)=0.23, p=0.638]). These results indicate that the retrieval of alcohol memories induces an elevation in corticosterone secretion.

### Stress exposure, corticosterone administration, and glucocorticoid receptor inhibition have no effect on alcohol-memory reconsolidation

Stress exposure after morphine memory retrieval was previously shown to impair the reconsolidation process ^55^. SAFit2, the FKBP51 inhibitor used here, has an anxiolytic effect, which emerges 16 hours after its injections ^56,65^. Therefore, it is possible that our finding that SAFit2 enhanced the reconsolidation and prevented the extinction of alcohol CPP 16 hours after its injection, can be explained by SAFit2’s anxiolytic properties. Therefore, we next tested whether exposure to stress can affect the reconsolidation of alcohol memories.

Mice were trained in the CPP procedure, as described above. Ten min following memory retrieval, mice were subjected to 30 min of restraint stress (see Methods), which was previously shown to induce activation of the HPA axis ^70^ and disrupt memory consolidation ^72–74^ and retrieval ^72,73,75^. Alcohol seeking was then monitored in four post-retrieval test sessions under extinction conditions (no alcohol) as above (Figure 6A) (see Experiment 7 in Methods).

**Figure 6.**
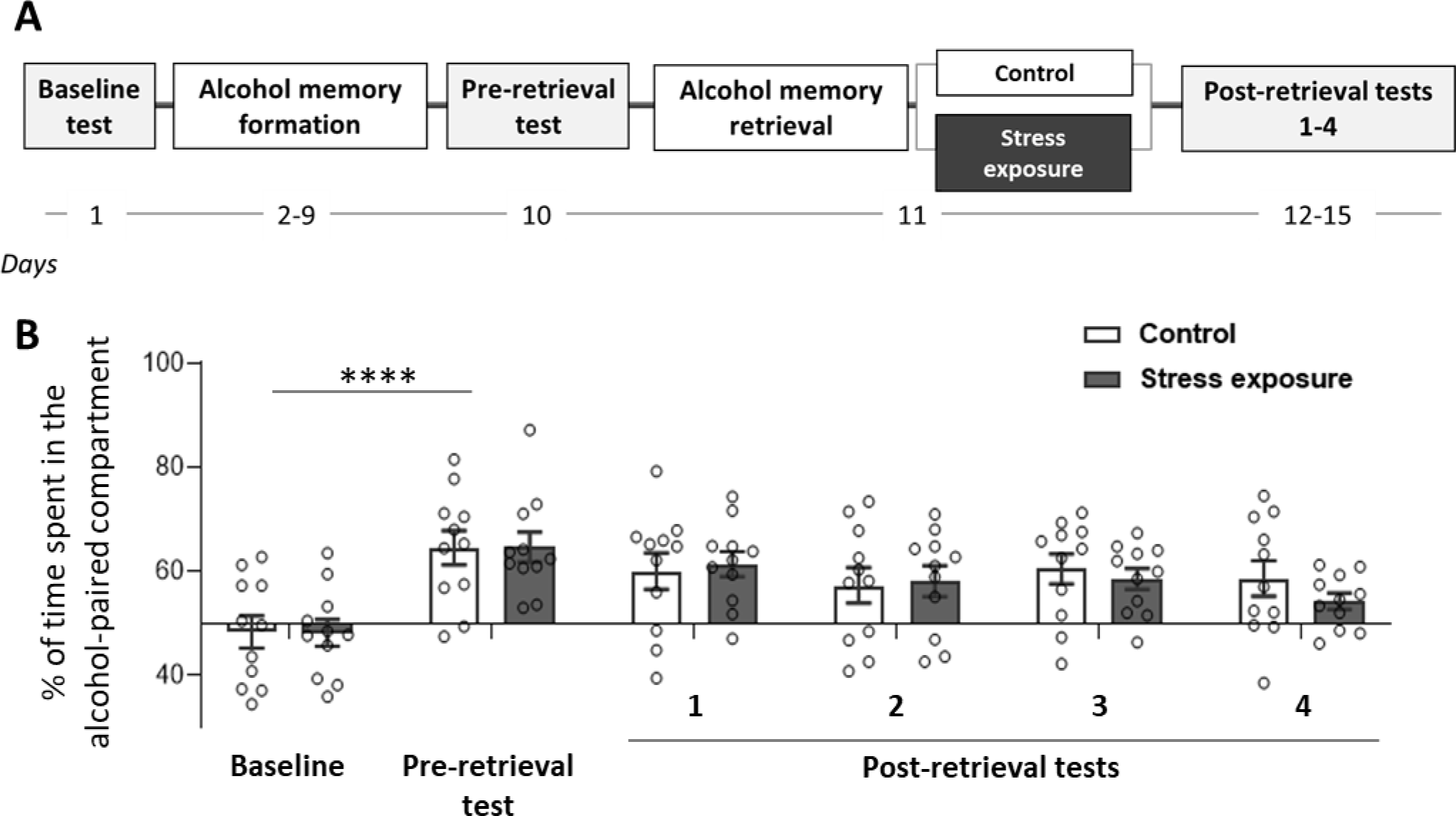
Stress exposure following alcohol-memory retrieval has no effect on alcohol-memory reconsolidation. **A.** Experimental design and timeline. Mice were trained in an alcohol CPP procedure. Ten min after the alcohol memory retrieval the mice were subjected to 30 min of restraint stress. The mice’s preference for the alcohol-paired compartment was monitored in four subsequential tests, 24 hours apart. **B.** Bar graphs indicate the mean percent of time spent in the alcohol-paired compartment ± SEM. n=11. **** p<0.0001 (Baseline test vs. Pre-retrieval test).

As expected, the mice displayed a higher preference for the alcohol-paired compartment after the conditioning phase, compared to their baseline preference (Figure 6B; t(21)=9.36, p<0.0001). There was no difference between the stress group and controls (mixed model ANOVA: no main effects of Group [F(1,20)=0.12, p=0.739] or Test [F(3,60)=1.54, p=0.214], and no Group X Test interaction [F(3,60)=0.26, p=0.538]).

Next, in two separate experiments, we examined the effect of systemic corticosterone administration and inhibition of the glucocorticoid receptors (GR) on memory reconsolidation (see Experiments 8 and 9 in Methods). Mice were trained in the CPP procedure as described above, and exhibited a preference for the alcohol-associated side in both experiments in the pre-retrieval test (Figure 7B – corticosterone: t(33)=11.7, p<0.0001; Figure 7C – mifepristone: t(24)=12.33, p<0.0001). Immediately after the memory retrieval session, we treated the mice with corticosterone (1 or 10 mg/kg, i.p., dose based on Wang et al. ^55^) (Experiment 8) or with the GR antagonist mifepristone (10 or 60 mg/kg, i.p., dose based on Lowery et al. ^76^) (Experiment 9), and control rats received corresponding vehicle. Then, we tested the preference for the alcohol-associated compartment over four test sessions under extinction conditions, as above.

**Figure 7.**
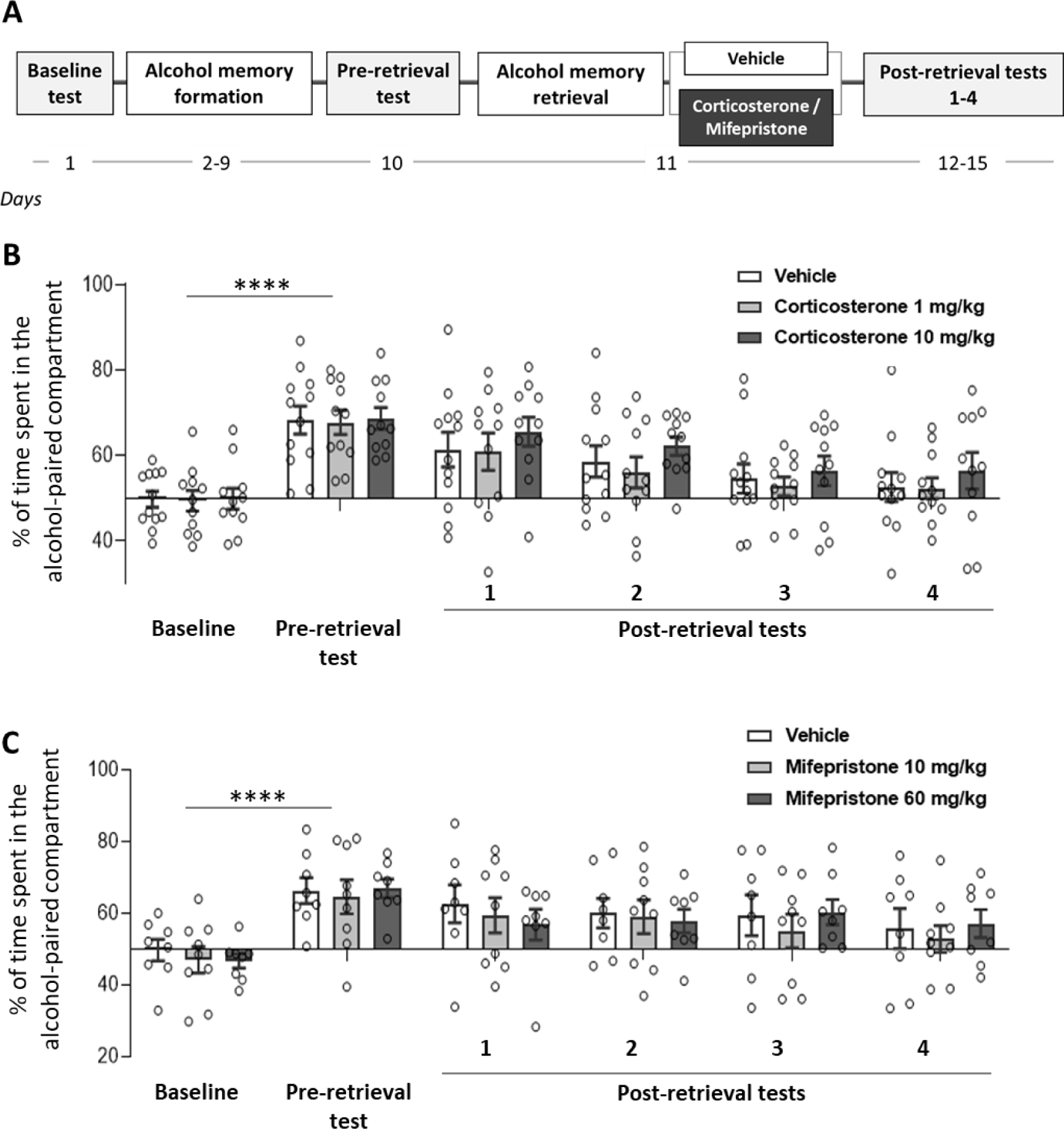
Administration of corticosterone or the glucocorticoid receptor antagonist mifepristone following alcohol-memory retrieval has no effect on alcohol-memory reconsolidation. **A.** Experimental design and timeline. Mice were trained in an alcohol CPP procedure. Immediately after the alcohol memory retrieval the mice were treated with vehicle, corticosterone (1 or 10 mg/kg), or Mifepristone (10 or 60 mg/kg). The mice’s preference for the alcohol-paired compartment was monitored in four subsequential tests, 24 hours apart. **B-C.** Bar graphs indicate the percent of time spent in the alcohol-paired compartment of the corticosterone-treated mice (B) and the mifepristone-treated mice (C). Bar graphs represent the mean±S.E.M. Corticosterone experiment: n=11-12; Mifepristone experiment: n=7-8. **** p<0.0001 (Baseline test vs. Pre-retrieval test).

We found that in both experiments, post-retrieval corticosterone (Figure 7B) or mifepristone (Figure 7C) treatment had no effect on the mice’s alcohol-seeking behavior throughout the post-retrieval tests (mixed model ANOVA, corticosterone: a main effect of Test [F(3,93)=11.93, p<0.0001], but no main effects of Treatment [F(2,31)=0.65, p=0.532], and no Group X Test interaction [F(6,93)=0.13, p=0.992]; mifepristone: no main effects of Test [F(3,66)=1.44, p=0.240], or Treatment [F(2,22)=0.14, p=0.870], and no Group X Test interaction [F(6,66)=0.60, p=0.732]).

These results suggest that exposure to external stressors, as well as to pharmacological manipulations on the activity of the HPA axis, has no effect on the reconsolidation of alcohol memories.

## 4. Discussion

We show that inhibition of FKBP51, a key regulator of the HPA axis, enhances the reconsolidation and prevents extinction of alcohol seeking. This effect was found selectively in female mice, in a place preference procedure, and is in line with our previous finding that alcohol memory retrieval in this procedure increases the *Fkbp5* mRNA expression in the mPFC ^25^. Moreover, we show that although the retrieval of alcohol memories in this procedure increases corticosterone levels in the blood, manipulations on the HPA axis activity following alcohol memory retrieval do not affect alcohol memory reconsolidation. Specifically, the application of stress, or systemic administration of corticosterone or the GR inhibitor mifepristone, shortly after alcohol memory retrieval did not affect alcohol seeking in the repeated extinction tests. We further show that while acute administration of alcohol leads to increased expression of *Fkbp5,* chronic-voluntary alcohol consumption has only minor and short-term effects on the expression of this gene. Finally, in contrast to our previous findings in a place conditioning reconsolidation paradigm with short-term alcohol exposure ^25^, the retrieval of alcohol memories using an odor-taste cue in rats with a history of long-term alcohol consumption, did not affect the *Fkbp5* expression. Together, these findings suggest that FKBP5/FKBP51 may play a role in the short and acute effects of alcohol and its memory reconsolidation, but not following long-term alcohol consumption.

### FKBP51 inhibition leads to persistent alcohol seeking in a sex-dependent manner

We have recently demonstrated in a mouse CPP procedure that alcohol memory retrieval increases the mRNA expression of *Fkbp5* ^25^. Here, we show that inhibition of FKBP51 by SAFit2 before the retrieval of alcohol memories prolongs the expression of alcohol place preference under conditions that induce extinction in controls, in females, but not in male mice. These findings can be interpreted as an enhancement of alcohol memory reconsolidation, and/or as attenuation of extinction. Given our previous findings of *Fkbp5* induction shortly after memory retrieval in this procedure ^25^, it is plausible that FKBP51, at least in females, is required for the flexibility of the memory. Accordingly, FKBP51 inhibition disrupts memory flexibility, resulting in persistent, inflexible behavior expressed as persistent alcohol-CPP and alcohol seeking.

We found that FKBP51 inhibition caused inflexible behavior only in females. Interestingly, the same FKBP51 inhibitor, SAFit2, was previously shown to attenuate stress-induced reinstatement of cocaine seeking in male, but not female rats ^77^, and to reduce alcohol drinking in male rats with stress history, whereas females persisted in drinking ^71^. Together, these studies suggest that the inhibitor affects drug-related behavioral flexibility in a sex-dependent manner, with males being more flexible than females. In line with this suggestion, a transgenic mouse model of *Fkbp5* overexpression displayed reduced pleasure-seeking behavior, as measured by sucrose preference test, in females but not in males ^78^. Thus, it seems that female mice with high FKBP51 changed their behavior compared to controls. An alternative interpretation of these findings may suggest that FKBP51 is a negative regulator of reward-seeking in females, and therefore its overexpression reduced sucrose preference and reward ^78^, whereas FKBP51 inhibition increased seeking for alcohol reward, as we show in our CPP experiment results.

The sex-dependent effects of FKBP51 manipulations may be linked to the interactions between FKBP51 and gonadal hormones. FKBP51 was reported to promote the activity of the androgen and estrogen receptors ^79,80^. In contrast, binding of FKBP51 to the progesterone receptor attenuates the receptor’s activity, whereas activation of the progesterone receptor induces the transcription of *FKBP5* ^81,82^. Indeed, the effectiveness of SAFit2 treatment in reducing stress-induced reinstatement of cocaine seeking was shown to be influenced by the rats’ estrous cycle phase at the time of the treatment ^77^. Moreover, using genetic tools, a thorough examination of two different conditional *Fkbp5* knockout mouse models revealed sex-dependent effects on behavior, brain structure, and gene expression profile ^83^. Our results further demonstrate that FKBP51-related manipulations can yield sex-dependent effects, also when applied in a non-stress-related setting.

Although the assessment of manipulations on memory reconsolidation typically requires post-retrieval treatments ^10,11^, we chose here to inject SAFit2 16 hours before the memory-retrieval session, as this inhibitor was shown to exert anxiolytic effects 16 hours post its systemic administration ^56,65,71^. Interestingly, in a previous study that was conducted solely on male mice, SAFit2 was shown to reduce the reinstatement of alcohol-seeking behavior in alcohol-CPP, when administered at a lower dose than the dose used in the current study two hours before the alcohol-primed reinstatement test, and in the absence of a retrieval procedure ^84^. Therefore, FKBP51 inhibition may have differential effects, depending on the timing and dose of administration. Moreover, as SAFit2 was administered 16 hours before the retrieval session, rather than after the retrieval procedure, it is possible that the treatment also affected the retrieval process and not only the post-retrieval reconsolidation process.

### Retrieval of alcohol memories via an odor-taste cue does not affect Fkbp5 expression in animals with a history of alcohol consumption

We recently reported an increased expression of *Fkbp5* after alcohol-memory retrieval using a contextual cue in a mouse CPP procedure ^25^, and indeed we show here that inhibition of FKBP51 affected the reconsolidation of contextual alcohol memories. In contrast, we found here that the retrieval of alcohol memories with an odor-taste cue, in rats with a history of long-term alcohol consumption, did not affect the mRNA expression of *Fkbp5*.

Importantly, based on previous studies, we used mice for the experiments that employed the CPP paradigm, and rats in the experiments that included memory retrieval using an odor-taste cue after IA2BC training ^20,25,64,85,86^. Therefore, the difference in the species possibly contributed to the discrepancy in the effects of each retrieval method on *Fkbp5* expression. Additionally, the memory-formation phase in the CPP procedure includes only four injections of alcohol, whereas, during the IA2BC training, the rats are given access to alcohol bottles for several weeks, allowing the formation of memories that are stronger and older at the time of the retrieval ^11,87^.

It is possible that prolonged alcohol exposure in the IA2BC procedure can potentially alter *Fkbp5* responsiveness to alcohol-related cues, as a history of heavy alcohol drinking was shown to attenuate cortisol reactivity to alcohol ^88,89^. In line with this possibility, a human study recently demonstrated blunted cortisol reactivity to alcohol-related cues in subjects who engage in binge drinking, compared with social drinkers ^90^.

Another methodological difference between the alcohol memory retrieval manipulation in CPP and the self-administration paradigms which can account for the different effects on *Fkbp5* expression is derived from the type of cues used for memory retrieval. Thus, the retrieval procedure in the CPP paradigm relies on contextual cues previously associated with the rewarding effect of the alcohol, which was given by injections ^91^. However, the retrieval procedure we used here after the long-term voluntary alcohol consumption utilizes the odor-taste properties of alcohol, which are intrinsic components of the oral alcohol consumption experience, and strongly associated with the reinforcing effect of alcohol ^20,86^. Therefore, the different nature of the cues that were associated with alcohol in each of the paradigms, and were later used to retrieve the alcohol memories, can also account for the different effects of each retrieval method on *Fkbp5* expression.

Interestingly, we also found that acute alcohol injection increased the mRNA brain expression of *Fkbp5,* in line with previous reports ^32–34^. We further show that this effect did not depend on the alcohol-drinking history of the mice, nor did we find significant effects of long-term alcohol consumption on this gene’s expression. These results indicate that although alcohol drinking triggers an immediate increase in *Fkbp5* expression, this alteration is not long-lasting and does not persist in the absence of alcohol. Together, our findings with both the direct alcohol exposure and the cue-induced alcohol memory retrieval, indicate that *Fkbp5* is more sensitive to acute manipulations related to alcohol than to long-term manipulations related to chronic alcohol exposure.

### Stress and glucocorticoid manipulations do not affect alcohol memory reconsolidation

Corticosterone promotes GR transcriptional activity, with *FKBP5* as one of its targets, and thereby acts as a positive regulator of *FKBP5* transcription ^35^. Therefore, it is possible that the elevation in *Fkbp5* expression we previously observed ^25^ was mediated by the increase in corticosterone secretion following memory retrieval. Indeed, we found that retrieval of alcohol memories led to increased levels of corticosterone in the plasma. Interestingly, previous studies have demonstrated that acute alcohol exposure increases corticosterone levels in the plasma and brain tissues ^92,93^, and that exposure to alcohol-associated cues facilitates this effect ^94^. Our present results indicate that the mere exposure to alcohol-related contextual cues can also increase corticosterone release. These results are in line with previous studies that demonstrated increased blood corticosterone levels, after re-exposure to a context that was associated with nicotine ^95^ or with cocaine ^53,54^.

Given our finding on increased corticosterone following alcohol memory retrieval, we speculated that stress and corticosterone elevation may play a role in alcohol memory reconsolidation. However, we found that restraint-stress exposure following alcohol-memory retrieval did not alter subsequent alcohol CPP. Similarly, we found that systemic administration of corticosterone or the GR antagonist, mifepristone, after memory retrieval, also failed to affect alcohol-memory reconsolidation. Our results differ from previous findings with other drugs, showing that morphine CPP in rats was attenuated by post-retrieval systemic injection of corticosterone or forced-swim stress, and the latter was reversed by intra-basolateral amygdala (BLA) mifepristone injection ^55^. Similarly, intra-BLA mifepristone injection following cocaine memory retrieval led to increased cocaine seeking, an effect interpreted as enhancement of memory reconsolidation ^54^. Moreover, indications for the ability of stress exposure and systemic or local pharmacological manipulations on the HPA axis activity, to impact the reconsolidation process were reported in multiple studies of fear memories ^51,52^ and non-drug appetitive memories ^96^. In light of contradicting findings in studies in the field of fear memories reconsolidation, it has been suggested that because of their function as consolidation enhancers, glucocorticoids are more efficient in the consolidation of new extinction memories than in the disruption of the reconsolidation of previously established memories ^52,97^. In the present study, the retrieval process, during which the mice were placed in the alcohol-associated context, lasted three min. In contrast, the retrieval sessions in the above-mentioned morphine ^55^ and cocaine ^54^ studies, lasted 10 and 15 min, respectively. Thus, the longer duration of the retrieval sessions in these studies had possibly increased the dominance of the extinction learning element over the reconsolidation element ^98^. Therefore, in these studies, unlike in the correct study, the cognitive process that took place during the retrieval session was more susceptible to HPA axis-related manipulations.

Interestingly, it was previously found that cortisol administration to human subjects, before exposure to alcohol-related images, reduced craving in patients with severe AUD, and enhanced craving in patients with less severe AUD ^99^. The history of alcohol consumption may therefore influence the effect of interventions related to the HPA axis on the retrieval or reconsolidation of alcohol memories.

Although in the present study we provide evidence for activation of the HPA axis following retrieval of contextual alcohol memories, the lack of effect of HPA axis-related manipulations on memory reconsolidation suggests that this system does not play a causal role in the reconsolidation of alcohol memories. Nonetheless, our finding that FKBP51 inhibition prior to alcohol memory retrieval led to a more prolonged and stable alcohol-seeking behavior in females, could be interpreted as the enhancement of reconsolidation or suppression of extinction. Thus, it is possible that the effect of FKBP51 inhibition was mediated by non-HPA axis-related mechanisms.

In summary, the results of the current study complement our previous observation that *Fkbp5* expression is increased after contextual alcohol memory retrieval ^25^, by demonstrating that FKBP51 inhibition leads to perseverative alcohol seeking in females when the memory is retrieved via its contextual cues. Taken together with previous findings, it is likely that FKBP51 plays a role in memory flexibility and changeability, in a sex-dependent manner. Interestingly, although we found that the retrieval of alcohol memories triggers activation of the HPA axis, this effect does not seem to play a causal role in alcohol memory reconsolidation, raising the possibility that FKBP51-related effects on memory flexibility are mediated via a non-HPA axis-related mechanism. Nonetheless, given previous findings implicating FKBP5/FKBP51 in AUD in humans ^46,49,100,101^, our present findings provide further evidence for the potential involvement of this protein in relapse to alcohol seeking.

## Acknowledgments

The research was supported by funds from the United States – Israel Binational Science Foundation (BSF) grant 2017022 and from the Israel Science Foundation (ISF) grant 508/20. Segev Barak is the Stephen Harper Chair of Translational Neuroscience at the Faculty of Social Sciences, Tel Aviv University.

## Author contributions

NR and SB designed the research; NR, CA, ML, KG and MLG performed the research; NR and SB analyzed the data and wrote the paper; FH and TH synthesized and provided the SAFit2 compound and reviewed the paper.

